# The selective 5-HT_1A_ receptor biased agonist, NLX-101, corrects anomalous behavioral phenotype in a mouse model of Fragile X syndrome

**DOI:** 10.1101/2025.09.16.676519

**Authors:** Ronan Depoortère, Michael R Tranfaglia, Adrian Newman-Tancredi

**Affiliations:** Neurolixis SAS, 2 rue Georges Charpak, 81100 Castres, France; FRAXA Research Foundation, Newburyport, MA 01950, USA

**Keywords:** Biased agonist, FMR1 KO2 mice, Fragile X syndrome, Mood/cognitive dysfunction, Neurodevelopmental disease, NLX-101, Serotonin 5-HT_1A_ receptors

## Abstract

Fragile X syndrome (FXS) is the most prevalent X-linked dominant autism spectrum disorder, causing a range of developmental problems, notably characterized by mild to severe mood/cognitive dysfunctions. NLX-101 is a highly selective and fully efficacious biased agonist at post-synaptic 5-HT_1A_ receptors, and has shown efficacy for reversal of sensory hypersensitivity and EEG anomalies in transgenic mouse models of FXS. Presently, we examined the ability of NLX-101 to normalize several aspects of behavioral anomalies displayed by adult male FMR1 KO2 mice, a transgenic murine model of FXS. FMR1 KO2 mice were treated with NLX-101 (0.64 & 2.5 mg/kg intraperitoneally) and tested sequentially in 1) the open-field test to study hyperactivity and stereotypies (self-grooming), 2) the three chamber partition test (social memory), 3) the nesting behavior test (daily living), 4) the novel object recognition test (working memory) and 5) the hyponeophagia (novelty suppression feeding) test (anxiety). Each test was separated by a three-day wash-out period. NLX-101 normalized hyperactivity and excessive self-grooming at both 0.64 and 2.5 mg/kg, whereas hyponeophagia, and deficits in working and social memory, were partially normalized at 0.64 mg/kg and fully at 2.5 mg/kg. Abnormal nest building was partially normalized at 2.5 mg/kg. In conclusion, NLX-101 exerts beneficial and dose-dependent activity against several behavioral and mood/cognitive deficits displayed by FMR1 KO2 mice. These results highlight the therapeutic potential of using a selective post-synaptic 5-HT_1A_ receptor biased agonist as a novel strategy to treat FXS, for which there is currently no approved efficacious and safe pharmacotherapy.

**SUMMARY:**

- Fragile X syndrome (FXS) is the most prevalent X-linked dominant autism spectrum disorder.
- NLX-101, a highly selective serotonin 5-HT_1A_ receptor agonist, was previously shown to reverse sensory hypersensitivity and brain electrical activity anomalies in a gene knock-out (FMR1 KO) mouse model of FXS.
- Here, NLX-101 corrected the anomalous behaviors (hyperactivity, stereotypies, anxiety, deficient social and working memories and nesting behavior) of FMR1 KO2 mice.
- Treatment with a selective 5-HT_1A_ receptor agonist such as NLX-101 could represent a promising strategy to treat cognitive and behavioral disturbance in FXS.

## INTRODUCTION

Fragile X Syndrome (FXS) is an inherited autosomal X-linked autism spectrum disorder (ASD) caused by lowered levels of the “Fragile X Messenger Ribonucleoprotein” (FMRP) (Dombrowski et al., 2002; Rousseau et al., 1995). FXS is a rare developmental disorder, with an estimated prevalence in the US of 1:7,000 for boys and 1:11,000 for girls (https://www.cdc.gov/fragile-x-syndrome/data/index.html) affecting the brains of both males and females, resulting in significant adverse effects on intellectual development and behavior, and constitutes the most common inherited form of intellectual disability. The expression of FMRP is under the control of the FMR1 gene, and the severity of symptoms/signs of FXS varies depending upon the degree of mutation of this gene. The FMRP is a repressor of translation and a key regulator of synaptic plasticity, and its absence has a profound impact on the neuronal circuitry and the normal functioning of the brain. As a consequence, patients with FXS can display a wide range of behavioral and mood/cognitive deficits, such as anxiety, hyperactivity, impulsivity, attentional problems, aggression, intellectual disabilities, repetitive actions, poor social interaction, speech and language delays, sensory hypersensitivity and propensity to epileptic seizures (Greiss Hess et al., 2016; Hagerman & Hagerman, 2002). Considering that there is no approved medication for the treatment of FXS, there is a great need for a novel, innovative, effective and safe pharmacotherapy.

At the preclinical level, drug candidates have often been evaluated using an FMR1 knockout (KO) mouse model first characterized by the Dutch-Belgian Fragile X Consortium (“Fmr1 knockout mice: a model to study fragile X mental retardation). This FMR1 KO mouse model was generated by a targeted insertion of a neomycin cassette into exon 5 of the FMR1 gene, resulting in a mouse that had low levels of FMRP and low levels of residual FMR1 mRNA. FMR1 knockout 2 (FMR1 KO2) mice were generated later by deletion of the promoter and first exon of FMR1 (Mientjes et al., 2006). The FMR1 KO2 construct lacks both the FMRP protein and FMR1 mRNA, and captures some of the behavioral abnormalities observed in human FXS patients, including hyperactivity, repetitive behavior, and deficits in learning and memory (Gaudissard et al., 2017). FMR1 KO2 mice have been used to identify potential drug candidates for treatment of FXS (Deacon et al., 2015; Tranfaglia et al., 2019)

The serotonin (5-hydroxytryptamine) 5-HT_1A_ receptor has recently emerged as a promising target for treatment of FXS. Thus, in a *Drosophila* model of FXS (homozygous dFMR1^Δ50^ mutants), treatment with eltoprazine, a 5-HT_1A_ receptor partial agonist, was found to ameliorate synaptic transmission, correct mitochondrial deficits, and ultimately improve motor behavior (Vannelli et al., 2024). Moreover, juvenile FMR1 KO mice had lower whole-brain 5-HT_1A_ receptor expression levels than age-matched controls, suggesting that a lack of sufficient 5-HT_1A_ receptor activation at a young age could impair proper neuronal development (Saraf et al., 2024).

In the present study we tested NLX-101 (a.k.a. F15599), a first-in-class, highly selective 5-HT_1A_ receptor biased agonist that preferentially targets post-synaptic receptors, particularly in cortical regions (Newman-Tancredi et al., 2022). NLX-101 shows rapid and robust activity in models of mood deficits, aggressivity and neuroplasticity (neuronal sprouting, increase BDNF levels, neurogenesis) (Cabanu et al., 2022; Vahid-Ansari et al., 2024; van Hagen et al., 2022) suggesting that it could be a promising drug candidate for treatment of ASD such as FXS. Accordingly, we recently showed that NLX-101 reduces audiogenic seizures (AGS) in developing FMR1 KO mice (Tao et al., 2023). In addition, NLX-101 extensively and potently increased survival of the mice over a similar dose range (1.2 to 2.4 mg/kg i.p.). Further, NLX-101 (1.8 mg/kg) was still strongly effective in reducing seizures even after repeated (daily) administration over 5 days, suggesting an absence of tachyphylaxis to the anti-seizure effects of the compound. Of note, NLX-112, a close chemical congener of NLX-101 which is also highly selective for 5-HT_1A_ receptors (Newman-Tancredi et al., 2022), similarly dose-dependently prevented AGS in juvenile FMR1 KO mice, an effect that was reversed by the selective 5-HT_1A_ receptor blocker WAY-100,635 (Saraf et al., 2024). In addition, NLX-101 improved auditory temporal processing in the same strain of FMR1 KO mice (Tao et al., 2025). Thus, whereas saline-treated FMR1 KO mice showed increased N1 (first negative peak in sound-evoked event related potentials) amplitudes and single trial power (STP), and reduced phase locking to auditory gap-in-noise stimuli versus wild-type mice, acute NLX-101 (1.8 mg/kg i.p.) significantly reduced STP at post-natal (PN) day 30. Intertrial phase clustering was also significantly increased by NLX-101 at PN21 and PN30, indicating improved temporal processing (Tao et al., 2025).

Taken together, these results suggest that NLX-101 could constitute a promising pharmacotherapeutic option to improve various dysfunctional aspects in FXS patients. In the present study, we characterized NLX-101 in adult male FMR1 KO2 (not FMR1 KO) mice using a battery of behavioral tests focused on hyperactivity, stereotypy, deficits in learning and memory and daily living functioning (Richter & Zhao, 2021). We used the open-field test for hyperactivity and stereotypies (self-grooming), the three chamber partition test for social memory, the nesting behavior test for daily living functioning, the novel object recognition test for working memory and the hyponeophagia (novelty suppressed feeding) test for anxiety.

## MATERIALS and METHODS

Experiments were carried out at the Institute of Ecology and Biodiversity Committee, University of Chile, Santiago, Chile.

### Animals

FMR1 KO2 male mice were provided by the FRAXA Research Foundation, MA, USA (https://www.fraxa.org/). FMR1 KO2 mice were generated by deleting the promoter and first exon of the FMR1 gene as described in (Mientjes et al., 2006), and were then backcrossed to a C57BL/6J background for more than eight generations. Similarly, wild-type (WT) mice were issued from a C57BL/6J background for more than eight generations. Initial stocks of mice were obtained from the Jackson Laboratory (Bar Harbor, ME, USA).

Mice were housed by four of the same genotype per cage in a temperature- and humidity-controlled (21 ± 1 °C, 60 ± 10% relative humidity) room with a 12 h light-dark cycle (lights on 7 a.m. to 7 p.m.). Mice were housed in commercial plastic cages (40 L × 23 W × 12 H cm^3^) with Aspen bedding on a ventilated rack system. Food and tap water were available *ad libitum*, except during behavioral tests.

Prior to behavioral testing, FMR1 KO2 and WT mice were randomly assigned to one of the treatment groups (vehicle, NLX-101 0.64 or 2.5 mg/kg i.p.), with the treatment maintained the same throughout the whole study for a given mouse.

On each experimental day, testing of all treatment groups was implemented in a balanced (i.e., similar number of animals per treatment group per day) and randomized manner (i.e., the order of testing of the animals determined using a randomization procedure).

Protocols were approved by the local ethical committee of the Institute of Ecology and Biodiversity Committee, University of Chile, Santiago, Chile and complied with the requirements of the UK Animals (Scientific Procedures) Act, 1986.

### Behavioral tests

Behavioral tests were implemented on mice 8 weeks of age (starting weight: 25.8 ± 0.6 g (mean ± SEM), range: 25–27 g; ending weight: 26.1 g ± 0.7 g (range: 25–28 g). in the following order, each test being separated from the previous one by 3 days:

1. Day 1: Locomotor activity and self-grooming in an open-field,
2. Day 5: Three chamber partition (social novelty)
3. Day 9: Nesting behaviour,
4. Day 13: Novel object recognition,
5. Day 17: Hyponeophagia (novelty suppressed feeding).

Each behavioral test was performed between 8 a.m. and 4 p.m., except for the nesting test which was run from 5 p.m. to 8 a.m.. Mice were dosed with vehicle or NLX-101 in the housing room prior to testing, and then brought to the experimental room to acclimate for 20 min before testing. Prior to each behavioral test, a mouse from a separate cohort that was not included in the study was placed in the experimental apparatus for 3 min. Then, this mouse was removed, and the apparatus was wiped with water-moistened and dry tissues before placing a study mouse into the apparatus. The aim was to create a low but constant background mouse odour for all experimental subjects.

#### Open-field (horizontal locomotor activity and self-grooming)

The open-field assay was performed using an automated system including activity monitor chambers (40 L × 40 W × 30 H cm^3^, Noldus, Wageningen, the Netherlands) with the associated EthoVision software (Noldus Information Technology Inc., Leesburg, VA, USA). Each mouse was first placed into a corner square facing the wall 60 min post-treatment, and was recorded for 30 min.

Horizontal locomotor activity was measured as the distance travelled in centimeters (cm); self-grooming was also recorded).

#### Three chamber partition social novelty

The apparatus was a rectangular three chambered box (Spectrum Diversified Designs, Inc., Streetsboro, OH, USA), each chamber measuring 40.5 L x 20 W × 22 H cm^3^. Dividing walls were made from clear Plexiglas, with small openings (10 W x 5 H cm^2^) that allowed access to each chamber, with the center chamber as the start location. A subject mouse was evaluated for its preference to explore a novel versus a familiar social stimulus mouse (from a separate cohort that was not included in the study). Exploration was defined as the time spent in the chamber with the novel mouse (NM) versus the time spent in the chamber with the familiar mouse (FM). During the exposure trial, the mouse was allowed to freely explore all three chambers for 10 min, with one mouse enclosed in a wire cage (3.8 cm diameter) in each of the side chambers. During the recognition trial, which took place immediately after the exposure trial, the subject mouse was allowed to freely explore all three chambers for 10 min, with one side chamber still containing the familiar mouse, while the other side chamber now containing a novel mouse. For each phase of the test, the amount of time spent in each chamber was recorded. An entry in a specific chamber was defined as all four paws in that chamber.

#### Nesting behavior

Nesting behavior was measured in a 40 L × 23 W ×12 H cm^3^ cage, with a single mouse per cage. Each cage was supplied 30 min (after the injection at 5 p.m.) with a “Nestlet,” a 5 cm square of pressed cotton batting (Ancare) available to arrange a nest. The test was run in the housing room. Scores of the quality of the nest were assessed the next morning at 8 a.m., when mice are naturally less active and typically sleep, using a 5 point scale the next morning, at the end of the dark phase:

1. The nestlet is largely untouched (>□90% intact),
2. The nestlet is partially torn up (50–90% remaining intact),
3. The nestlet is mostly shredded but often there is no identifiable nest site:□<□50% of the nestlet remains intact but□<□90% is within a quarter of the cage floor area, i.e., the cotton is not gathered into a nest but spread around the cage,
4. An identifiable, but flat nest:□>□90% of the nestlet is torn up, the material is gathered into a nest within a quarter of the cage floor area, but the nest is flat, with walls higher than mouse body height (curled up on its side) on less than 50% of its circumference,
5. A (near) perfect nest:□>□90% of the nestlet is torn up, the nest is a crater, with walls higher than the mouse body height on more than 50% of its circumference.

#### Novel object recognition

The novel object recognition apparatus was a Plexiglas box (26 cm length × 20 cm width × 16 cm height). Mice were first habituated individually to the experimental environment by allowing them to freely explore the box, which was empty, for 20 min per day for two consecutive days before testing. The test involved two consecutive trials, each 5 min in duration separated by an interval of 30 min. In the exposure trial, two identical (i.e., familiar) objects (4 cm diameter × 2 cm H, made of plastic) were placed in the center of the chamber, 8 cm apart, and the mouse was allowed to freely explore the objects for 5 min. For the recognition trial, one familiar object (FO) was replaced with one novel object (NO), with the mouse again left for 5 min of exploration. Object exploration was defined as the mouse sniffing or touching the object with its nose, vibrissa, mouth, or forepaws. The time spent near or standing on top of the objects without interacting with the object was not counted as exploration.

#### Hyponeophagia (novelty suppressed feeding)

Mice were food restricted overnight and tested the next morning; twenty minutes prior to the test, each mouse was placed into a temporary holding cage to prevent social transmission of food preferences. Testing was conducted in a white Perspex chamber (30 L × 30 W × 5 H cm^3^) with three white walls and a fourth wall of transparent plastic to allow for observation of the mouse. A food well (1.2 cm diameter, 0.9 cm height) was glued to the base in the center of the test chamber. Each mouse was placed into the chamber facing away from the food well containing Nestle Carnation sweetened condensed milk diluted 50:50 with water. The latency time from placement in the test chamber to the start of a proper drinking bout, defined as drinking continuously was measured with a stopwatch with a precision of 0.1 s.

#### Pharmacological compounds

NLX-101 (a.k.a. F15599: N-{[1-(3-chloro-4-fluorobenzoyl)-4-fluoropiperidin-4-yl]methyl}-N-[(5-methylpyrimidin-2-yl)methyl]amine; hemi-fumarate salt) was provided by Neurolixis, dissolved in distilled water and administered i.p. at a volume of 5 ml/kg body weight; doses correspond to the weight of the base. Vehicle consisted of distilled water. NLX-101 or vehicle administration was performed 60 min prior to test (recognition trial for the Three chamber partition social recognition and the Novel object recognition tests). For the nesting behavior experiment, drug treatment was administered in the evening right before overnight recording. Experimenters were blind to the mouse genotype and treatment condition throughout all behavioral tests and data analysis. The doses of NLX-101 were chosen based on efficacy data against audiogenic seizures (Tao et al., 2023) or auditory temporal processing (Tao et al., 2025; Tao et al., 2023) in FMR1 KO mice.

#### Data analysis

Data were analyzed with one-way ANOVAs, followed by Holm-Sidak’s post-hoc tests wherever appropriate, and results are reported in the figure legends (ANOVAs) and by statistical symbols in the figures. Statistical analyses were implemented with the Prism^®^ V10.4.1 software (GraphPad Software, Boston, MA, USA).

## RESULTS

### NLX-101 normalizes several aspects of abnormal behavioral phenotype of *FMR1* KO2 mice

In the Open-field test, vehicle-treated WT mice travelled, on average, 3335 cm (1^st^ set of bars, upper left panel, Fig. 1); In contrast, vehicle-treated *FMR1* KO2 mice displayed hyperactivity, and travelled 2 ½ times more (8162 cm; 2^nd^ set of bars). Administration of NLX-101 (0.64 or 2.5 mg/kg i.p.) nearly fully reversed hyperactivity in KO2 mice, with an averaged distance travelled comparable to that of WT mice at 2.5 mg/kg.

**Figure 1:**
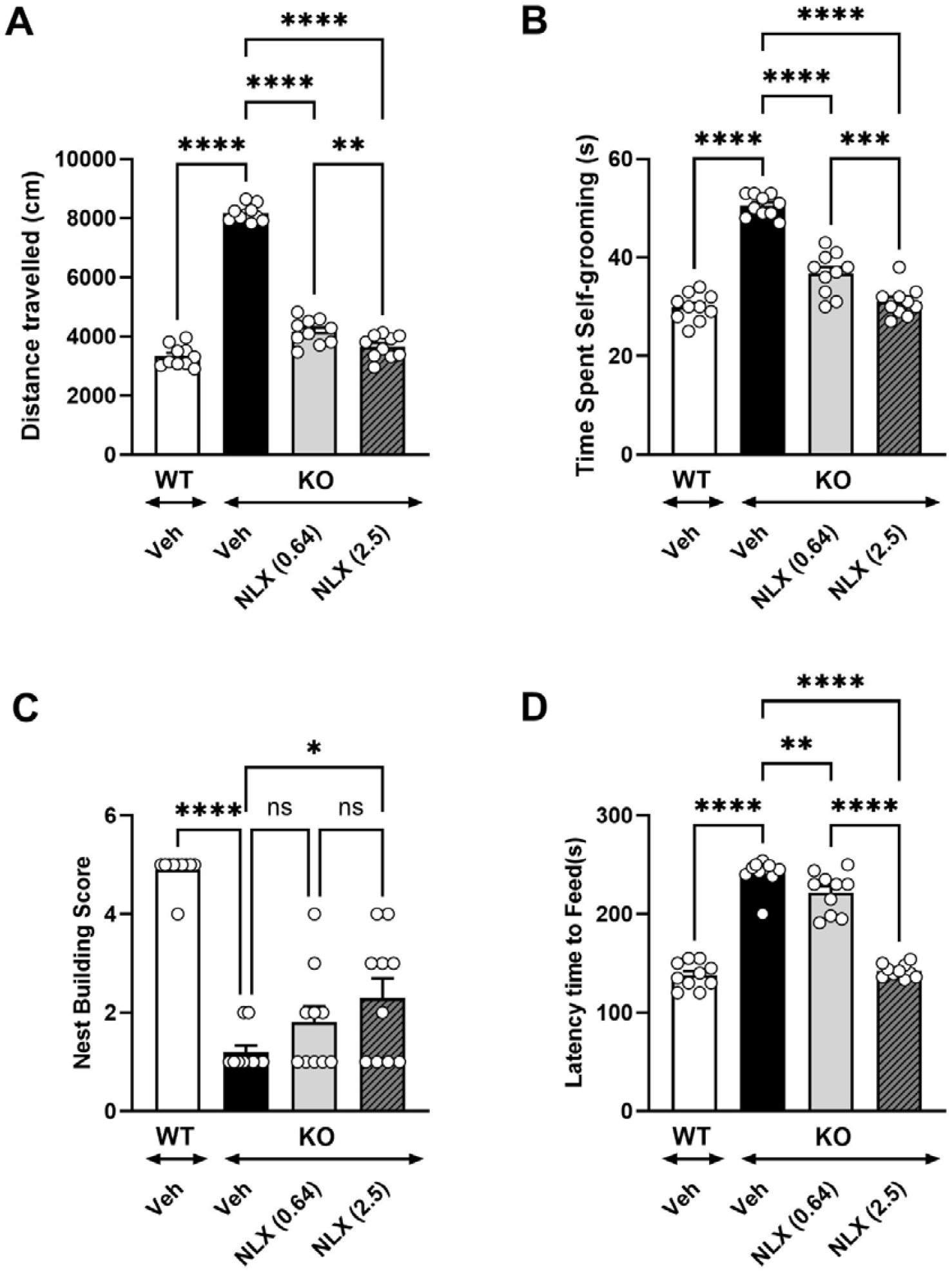
Panel A: hyperactivity in the open-field test; Panel B: self-grooming in the open-field test; Panel C: nest building test; Panel D: hyponeophagia in the novelty-suppressed feeding test. Bars are mean with SEM, symbols are individual data. N=8 to 10 mice per group. KO: knock-out, Veh: vehicle, WT: wild-type; numbers in parentheses are doses in mg/kg i.p.. * p<0.05, ** p<0.01, **** p<0.0001, ns: not significant, Holm-Sidak’s post-hoc test following significant one-way ANOVA. For hyperactivity: F (3,36) = 377.3, for self-grooming: F (3,36) = 36.5; for hyponeophagia: F (3,36) = 128.2; for nest building: (3,36) = 388.7; all P’s<0.0001.

Similarly, *FMR1* KO2 mice spent 69% more time self-grooming than their WT counterparts (29.9 s vs 50.5 s: compare 1^st^ and 2^nd^ bars, upper right panel); NLX-101 greatly and significantly diminished self-grooming at both doses tested.

In the Nest building test, WT mice attained on average a near-maximal score (4.9 out of 5), whereas *FMR1* KO2 mice scored only 1.2, i.e., the nestlet was left largely untouched. NLX-101 dose-dependently and significantly (at 2.5 mg/kg) augmented the nest building score.

Lastly, in the Hyponeophagia test, whereas the latency time for WT mice to engage in feeding was 138.0 s, the latency time for FMR1 KO2 mice was almost twice as much (241.2 s: lower left panel). This increase in latency time was fully and significantly reversed with the higher dose of NLX-101 (2.5 mg/kg).

### NLX-101 attenuates working and social memory deficits in FMR1 KO2 mice

In the Novel Object Recognition test, vehicle-treated WT mice spent more than twice as much time exploring the novel (unfamiliar) versus the familiar object (6.5 vs 14.1 s: compare 1^st^ and 2^nd^ bars, panel A, Figure 2). In contrast, vehicle-treated FMR1 KO2 mice spent less time exploring the novel object than the familiar object. Administration of 0.64 mg/kg NLX-101 tended to increase the time spent by FMR1 KO2 mice exploring the novel object, and the dose of 2.5 mg/kg increased it significantly.

**Figure 2:**
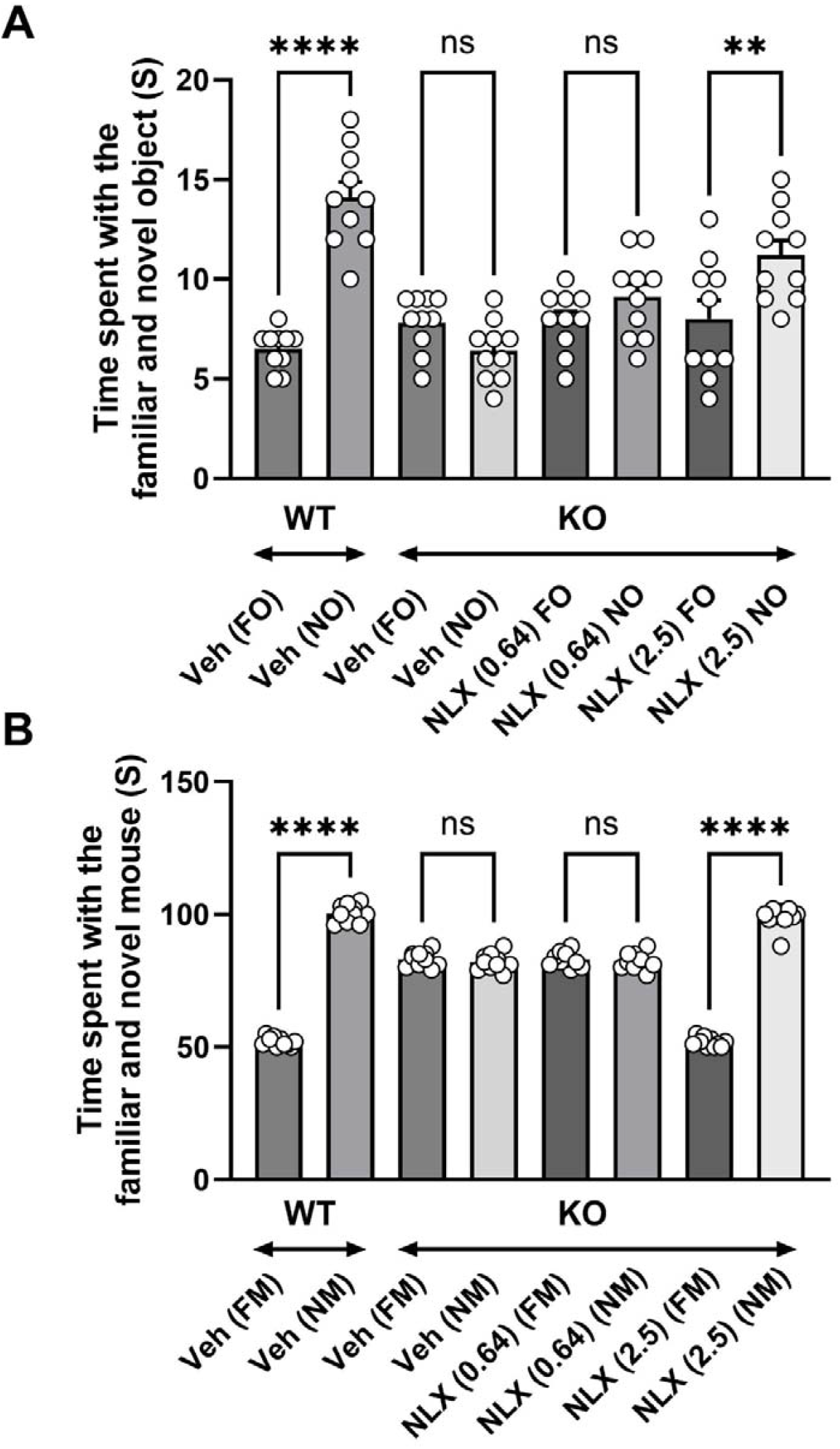
Panel A: working memory in the novel object recognition test. Panel B: social memory in the three-chamber partition test). Bars are mean with SEM, symbols are individual data. N=10 mice per group. FM: familiar mouse, KO: knock-out NO: novel mouse, WT: wild-type; numbers in parentheses are doses in mg/kg i.p.. ** p<0.01, **** p<0.0001, ns: not significant, Holm-Sidak’s post-hoc test following significant one-way ANOVA. For novel object recognition: F (7,72) = 377.3; for social interaction: F (7,72) = 396.6; both P’s<0.0001.

In the Three chamber partition test, vehicle-treated WT mice spent twice as much time interacting with the novel (unfamiliar) mouse than with the familiar mouse (52.0 s vs 100.3 s: compare 1^st^ and 2^nd^ bars, panel B, Figure 2). Vehicle-treated FMR1 KO2 mice spent a similar amount of time interacting with the familiar or novel mouse. Whereas 0.64 mg/kg NLX-101 was ineffective in increasing the time spent interacting with the novel mouse, 2.5 mg/kg significantly and fully restored the time ratio of exploring the novel versus the familiar mouse, up to the level seen in the WT vehicle-treated cohort.

Figure 2 about here

## DISCUSSION

The main finding of the present study is that the selective 5-HT_1A_ receptor biased agonist NLX-101, when administered i.p. under repeated/acute conditions (5 injections, each separated by 3 days of wash-out), almost completely reversed the anomalous behavioral phenotype of adult male FMR1 KO2 mice, a transgenic murine model of FXS. Thus, NLX-101 completely normalized (i) hyperactivity and excessive self-grooming in the open-field test, (ii) hyponeophagia in the novelty suppressed feeding (NSF) test, and (iii) social memory in the three-chamber partition test. In addition, (iv) NLX-101 attenuated the mice’ deficit of working memory in the NOR test and their abnormal nest building behavior.

### NLX-101 improves abnormal behavioral measures in FMR1 KO2 mice

This series of positive data using a battery of behavioral tests in FMR1 KO2 mice strengthens previous positive results obtained with the FMR1 KO murine transgenic construct (see Introduction). As a reminder, NLX-101 reduced AGS and increased survival of juvenile mice subjected to loud noise (Tao et al., 2023), and also normalized anomalous EEG phenotypes in developing FMR1 KO mice (Tao et al., 2025). Together with the present observations, these cumulative data indicate that the beneficial activity of NLX-101 on AGS and anomalous EEG profile translates to improvement in the abnormal behavioral phenotype of the FXS mouse model. Of note, the doses at which NLX-101 was active in the present battery of behavioral tests in FMR1 KO2 mice are consistent with those in which it was active against AGS and abnormal EEG phenotype in FMR1 KO mice. Thus, both murine constructs appear to respond similarly to the activity of NLX-101.

NLX-101 dose-dependently countered the anomalous hyperactivity, excessive self-grooming and hyponeophagia behaviors. NLX-101, at the higher dose, also corrected deficits in nest building, novel object recognition and social recognition, indicating a broad activity of the compound against a wide range of behavioral impairments in FMR1 KO2 mice, and suggesting that it has promising features for potential treatment of FXS in humans. In the nest building test, NLX-101 appeared to be less efficacious at restoring performance of FMR1 KO2 mice to levels close to those of control mice. This may reflect region-specific effects of the compound and the distinct neural circuits underlying each behavioral domain. NLX-101 is a selective 5-HT_1A_ recptor agonist that preferentially targets cortical receptors, particularly in the medial prefrontal cortex (mPFC), with minimal activation of 5-HT autoreceptors in the raphe nuclei. Nesting behavior is considered an ethologically relevant measure of species-typical behavior and motivation, and involves broader and more integrative neural substrates. It depends on the medial preoptic area, hippocampus, and possibly subcortical networks regulating motivation, arousal, and thermoregulation (Deacon, 2006). These regions may be less responsive to the cortical-selective action of NLX-101, or the serotonergic modulation in these circuits may not be sufficient to normalize nesting behavior in FMR1 KO2 mice. This could be explored by local administration of the compound in different brain regions, as has already been done in some studies in mice and rats (Papp et al., 2024; Stein et al., 2013).

### Behavioral improvement and neuroplastogenic properties of NLX-101

It should be noted that there was a 3-day wash-out period between each of the behavioral tests, in order to avoid NLX-101 dosing carry-over effects which might have confounded results. This duration of wash-out should be amply sufficient to avoid pharmacokinetic interactions, given that the half-life of NLX-101 in C57BL/6J mice is about 2 h in plasma, and 2.8 h in whole brain (Monteiro-Fernandes et al., 2025), so NLX-101 exposure after the 3-day wash-out would therefore be negligible. However, it cannot be discounted that the striking neuroplastogenic effects of NLX-101 on serotonin neurons (i.e., rapid and sustained changes after a single administration) (Vahid-Ansari et al., 2024) and its induction of neurogenesis and BDNF release (Cabanu et al., 2022; van Hagen et al., 2022) could have produced changes which may have influenced some of the effects observed in the later behavioral tests. This point would, ideally, require additional testing of NLX-101 in separate groups of animals for each test, but such a study has marked ethical and cost implications by multiplying the number of mice to be tested. On a therapeutic level, however, a putative neuroplastic effect of NLX-101 may be associated with a desirable disease-modifying activity which would be highly beneficial for individuals with FXS. Such a hypothesis would require appropriate clinical investigation in FXS patients, an objective which is facilitated by the fact that NLX-101 has already safely undergone a single-dose Phase 1 trial in adult males, has an open IND (Investigational New Drug) authorization in the USA and has been granted Orphan Drug Designations for treatment of FXS in both the USA and the European Union.

An important behavioral trait of FXS which was not investigated in the present study is aggressivity (Tranfaglia, 2011), a disturbance which is usually treated off-label with multi-target antipsychotics such as aripiprazole or risperidone (Protic et al., 2022). Of note, NLX-101 robustly reduced aggressive behavior (frequency of attack bites and sideways threats) when microinjected into the ventral orbital prefrontal cortex of CF-1 male mice (Stein et al., 2013). The anti-aggressive activity of NLX-101 was prevented by intra-cortical microinjection of the selective 5-HT_1A_ receptor antagonist WAY100,635, demonstrating the receptor specificity of NLX-101. These observations support the notion that NLX-101 acts principally through targeting of cortical 5-HT_1A_ receptors and suggest that, in addition to improving mood and cognition, it could also dampen aggressive behavior in subjects with FXS.

### Limitations and future directions

Looking ahead to further investigation, several points should be noted. Firstly, the present study used only male FMR1 KO2 mice, and it would be informative to determine the effects of NLX-101 in female mice. Secondly, it would be interesting to test the effects of the selective 5-HT_1A_ receptor antagonist, WAY-100,635 in FMR1 KO2 mice to assess if there is a basal 5-HT_1A_ tone that provides some level of behavioral protection. Moreover, combining WAY-100,635 with NLX-101 would provide confirmation that the beneficial behavioral effects of the latter stem from activation of 5-HT_1A_ receptor, consistent with the exceptional 5-HT_1A_ receptor selectivity of the compound. Indeed, whenever WAY-100,635 was previously tested in combination with NLX-101 in various rodent (rats, mice) behavioral models, it systematically blocked the effects of the latter (see (Newman-Tancredi et al., 2022) for detailed review). Lastly, NLX-101 was tested in FMR1 KO2 mice aged from 8 to 11 weeks, but it would be informative to investigate the effects of NLX101 in even younger mice (although they may be too immature to carry out some of the behavioral tests, notably nest building).

## Conclusions

The present data support the potential benefit of selective serotonin 5-HT_1A_ receptor activation as a treatment for FXS. The behavioral improvement elicited by NLX-101 in FMR1 KO2 mice is consistent with that of previous studies in FMR1 KO mice (reduced auditory hypersensitivity and anomalous EEG profile) and with rodent studies showing improved performance in models of mood and cognition. Nevertheless, demonstration of clinical therapeutic efficacy of NLX-101 is necessary to support further development of the compound for treatment of the neuropsychiatric and neurological signs/symptoms of people with FXS and, potentially, other ASD patients.

## ACKNOWLEDGEMENT

We wish to thank Pablo Cornejo and Robert Deacon for carrying out the experimental work.

